# Health assessment of the pink land iguana, *Conolophus marthae*

**DOI:** 10.1101/2021.08.26.457766

**Authors:** Gabriele Gentile, Giuliano Colosimo, Carlos A. Vera, Glenn P. Gerber, Hans Westermeyer, Christian Sevilla, Gregory A. Lewbart

## Abstract

The pink land iguana, *Conolophus marthae*, is one of four species of iguanas (three terrestrial and one marine) in the Galápagos Islands, and the only one listed as critically endangered by the IUCN. The species can only be found on the north-west slopes of the highest volcano on the island of Isabela and was first described to science in 2009. As part of a population telemetry study, a health assessment was authorized by the Galápagos National Park. Wild adult iguanas were captured on Wolf Volcano in September 2019 and April 2021 to record morphological and physiological parameters including body temperature, heart rate, body measurements, intraocular pressures, tear formation, and infrared iris images. Blood samples were also collected and analyzed. An i-STAT portable blood analyzer was used to obtain values for base excess in the extracellular fluid compartment (BEECF), glucose, hematocrit, hemoglobin, ionized calcium (iCa), partial pressure of carbon dioxide (pCO_2_), partial pressure of oxygen (pO_2_), percent oxygen saturation (sO_2_%), pH, potassium (K), and sodium (Na). Standard laboratory hematology techniques were employed for packed-cell-volume (PCV) determination. When possible, data were compared to previously published and available data for the other Galápagos iguanas. The results reported here provide baseline values that may be useful in detecting changes in health status among pink land iguanas affected by climate change, invasive species, anthropogenic threats, or natural disturbances.

## Introduction

The Galápagos pink land iguana, *Conolophus marthae,* was first sighted on Wolf Volcano (WV hereafter), Isabela Island, in 1986, but was not described to science as a separate and unique species until 2009 (Gentile & Snell, 2009). The species is under the protection of Galápagos National Park Directorate (GNPD) and listed in Appendix II of the Convention on International Trade in Endangered Species of Wild Fauna and Flora (CITES, Appendices I, II and III, 2021). It is classified as critically endangered by the IUCN Red List of Threatened Species (Gentile, 2012). Among the major threats for the species we recognize: a single and very small population (200 < N < 300 adult individuals); small area of distribution estimated at no more than 25 km^2^; lack of recruitment, with no hatchlings and very few juveniles observed since 2005; introduced alien species such as cats and rats; and volcanic eruption (WV is an active volcano that last erupted in 2015). With such a small population, these animals are at risk of extinction, from both natural and anthropogenic events. Although information on the natural history and biology of *C. marthae* has been accumulating since its description to science (Gentile & Snell, 2009; Gentile *et al.*, 2009; Fabiani *et al.*, 2011; Gentile, 2012; Gentile *et al.*, 2016; Di Giambattista *et al.*, 2018; Colosimo *et al.*, 2020), very little is known about the species’ overall health and baseline medical parameters. This called for urgent action aimed at providing data about the health status of this flagship species.

Peripheral blood biochemical, blood gas, and hematology parameters are useful for assessing lizard health (Geffre *et al.*, 2009; Klaphake *et al.*, 2017; Arguedas *et al.*, 2018; Lewbart *et al.*, 2019). It is important to establish and publish species-specific baseline physiological parameters for healthy individuals since disease, injury, pollutants, starvation and other factors can result in blood value perturbations. Several reference intervals have been established for iguanids and include: *Conolophus pallidus* and *C. subcristatus* (Lewbart *et al.*, 2019), *Amblyrhynchus cristatus* (Lewbart *et al.*, 2015), *Basiliscus plumifrons* (Dallwig *et al.*, 2010), *Cyclura cychlura inornata* (James *et al.*, 2006), *Cyclura ricordii* (Maria *et al.*, 2007), and *Iguana iguana* (Dennis *et al.*, 2001; Divers *et al.*, 1996; Harr *et al.*, 2001; Hernandez-Divers *et al.*, 2005; Wagner & Wetzel, 1999).

As part of a remote monitoring population study, a health assessment study for the pink iguanas was authorized by the GNPD. A detailed description of the satellite tracking study is beyond the scopes of this article, and we refer to Loreti and colleagues (2019 and 2020) for further information. Nevertheless, we took advantage of this study and proceeded in two ways: *i)* we sampled wild pink iguanas on WV, Isabela Island, in September 2019 and April 2021 to analyze blood chemistry and hematological parameters and establish an intra-specific baseline report of such parameters; *ii)* we collected all published blood chemistry data available for other Galápagos iguanas to perform, when possible, an inter-specific comparison of such parameters. For wild samples collected in 2019 and 2021 a complete veterinary health examination was performed. The examination included measurement of body temperature, heart rate, length, body weight, intraocular pressure, ocular tear production, collection of blood samples, ectoparasites, and, in some cases, feces. All procedures were in accordance with the ethics and animal handling protocols approved by the GNPD. In the current study, we report on the blood chemical analysis and status of clinically healthy wild adult Galápagos pink land iguanas.

## Materials and Methods

### Ethic statement

This population health assessment was conducted on Wolf Volcano, Isabela Island, Galápagos archipelago (Ecuador), and was authorized by the Galápagos National Park Directorate (Permit # PC-04-21 issued to G. Gentile). The techniques used during this health assessment were also approved by the University of Rome “Tor Vergata” ethics and animal handling protocol and the San Diego Zoo Wildlife Alliance IACUC. All procedures were carried out by a licensed veterinarian and author of this study (GL).

### Iguana capture, measurement, and sampling

We captured, tagged, and sampled 15 and 27 adult pink iguana individuals during field expeditions conducted in September 2019 and April 2021, respectively. Individuals were captured in an area on WV comprising approximately 6 km^2^ at an altitude of 1600-1700 m. All iguanas were captured by hand or noose, either inside of or adjacent to their burrows. The animals were quickly transported to a field laboratory area (usually located within 10 meters of the capture site) for blood drawing, usually within 5 minutes from capture. A blood sample of approximately 2.5 mL was obtained from the coccygeal hemal arch of each iguana using a heparinized 22-gauge 3.8 cm needle attached to a 10.0 mL syringe. The blood was divided into sub-samples and stored in a field cooler kept refrigerated using icepacks. Once the blood sample was secured the animal was examined, measured, weighed, and a custom designed GPS tracker was attached (Loreti *et al.* 2019, 2020 for further details on the tracking devices).

Cloacal temperature and heart rate were recorded shortly after the blood was taken, and usually within 5 minutes of capture. Heart rate was recorded via a Doppler ultrasound probe (Parks Medical Electronics, Inc., Aloha, Oregon, USA) over the heart. An EBRO^®^ Compact J/K/T/E thermocouple thermometer (model EW-91219-40; Cole-Parmer, Vernon Hills, Illinois, USA 60061) with a T-PVC epoxy-tipped 24 GA probe was used to determine core body temperature. Snout-vent length (SVL) and tail length (TL) were recorded with a flexible measuring tape and used to determine the total length. Body mass was measured with a digital scale (± 0.1 kg). The sex of the iguanas was determined by the presence or absence of hemipenes or by visually inspecting secondary sexual characteristics (presence/absence of femoral pores, size of the individual, prominence of nuchal crest). Prior to release, ectoparasite load was determined by counting the number of ticks as in Onorati and colleagues (2016). For each animal, a sample of ticks was collected and preserved in 70% ethanol. In 2021 close inspection of all animals revealed populations of small red mites associated with the skin over the cranial trunk and shoulder areas. Samples were collected and preserved in 70% ethanol. Each animal was also scanned for the presence of a Passive-Integrated-Transponder (PIT) tag and checked for the presence of a brand. For never-before captured individuals a PIT tag was placed under the femoral skin of the right leg for long term identification and population monitoring.

A complete examination of the eyes was performed by a single evaluator (GAL). Within 15 minutes of capture, internal ocular pressure (IOP) measurements were taken of the left (IOP_L) and right (IOP_R) eye using a rebound tonometer (TonoVet^®^, iCare, Tiolat, Helsinki, Finland). Intraocular pressures were measured on the Tonovet’s^®^ rebound tonometer on undefined patient setting (p). Disposable probes were used and changed between every patient. The tonometer was held in position perpendicular to the patient eye and approximately 5 mm from the corneal surface. Tear production was measured using the endodontic absorbent point paper test (EPPT) in all 15 iguanas sampled in 2019 but not for samples collected in 2021. Size 30, 40, and 45 endodontic paper points (Parallax^®^ Veterinary Absorbent Paper Points) were used. The tapered end of the clean points was placed into the fornix in both the left and right eye. The paper was removed after 60 seconds and the length of moisture that was wicked on the point was measured with a millimeter ruler. External infrared photographs of each eye (Panasonic^©^ Lumix DMC-ZS50 12.1MP) were obtained for each animal sampled in 2019 and reviewed for ocular abnormalities by a board-certified veterinary ophthalmologist (HW). Before releasing each animal in the exact location where it was caught, a unique ID number was painted on both flanks and the tip of the tail (to prevent recapture) using a non-toxic, washable white paint.

### Hematology parameters

The blood samples were used for measuring various hematological parameters: approx. one drop was used for lactate analysis; approx. one drop for glucose analysis; about 0.1 mL was loaded into an i-STAT Clinical Analyzer (Heska Corporation, Fort Collins, Colorado, USA) utilizing Chem8 cartridges (see later in text for details); about 0.05 mL was used for centrifugation with a portable microcentrifuge (Eppendorf North America, Inc., centrifuge model 5424, 5 min. at 14,000 G) to determine packed cell volume (PCV) and total solids (TS). The PCV was determined by measuring the percentage of cellular material compared to plasma in the tubes. Two drops of plasma were placed on a refractometer (Ade Advanced Optics, Oregon City, Oregon 97045, USA) and the total solids values recorded. We additionally used approx. one drop of blood for making blood films on clean glass microscope slides (samples not yet analyzed).

The i-STAT Clinical Analyzer is a handheld, battery-powered, device that measures selected blood gas, biochemical, and hematology parameters using approx. 0.095 mL of non-coagulated whole blood. The following parameters were measured: base excess in the extracellular fluid compartment (BEECF), bicarbonate (HCO_3_^−^), glucose, hematocrit, hemoglobin, ionized calcium (iCa), partial pressure of carbon dioxide (pCO_2_), total carbon dioxide (TCO_2_), partial pressure of oxygen (pO_2_), pH, potassium (K), and sodium (Na). The i-STAT automatically produces temperature corrected values for pCO_2_, pH and pO_2_ once the animal’s body temperature is entered. Blood lactate was determined using a portable Lactate Plus™ analyzer (Nova Biomedical, Waltham, Massachusetts, 02454 USA). A glucometer (Accu-Check^®^ Active, Roche) was sometimes used to obtain near instant glucose values in the field and compare them to the glucose values obtained by the i-STAT Clinical Analyzer.

### Data compiling procedure

We reviewed available and published health and hematological data from other Galápagos iguanas. Data were sourced primarily from two publications by Lewbart and colleagues (2015, 2019) from individuals of *C. pallidus*, *C. subcristatus*, the two other species of Galápagos land iguanas, and *A. cristatus*, the Galápagos marine iguana. The protocols used while sampling individuals of these other species are the same as those adopted for this study, in fact they were performed by the same author (GL). For this reason, when possible, the data were used to perform inter-specific comparisons.

### Data analysis

Unless otherwise stated, all statistical analyses and data manipulations were performed in R v4.0.5 (R Core Team 2021). We compared morphological, blood gas, and hematology parameters between pink iguana samples and, when possible, we compared these results to parameters available for other species of Galápagos iguanas. We divided the analytical procedure in two parts: intra-specific and inter-specific.

### Intra-specific

We first built boxplots using SVL and mass from each individual. After removing potential outliers we calculated summary statistics (mean, standard deviation, minimum and maximum values) for these two variables while accounting for sex differences.

In 2019 we collected some hematological parameters using alternative protocols to allow a comparison between alternative methodologies. For example, we used two different i-Stat cartridge types (Chem8 and CG-8), and in some cases we also repeated the measurement using standard lab techniques and the use of a micro-centrifuge (see description above). This procedure allowed us to compare whether data collected using different methodologies were significantly different from each other, and we report on this below. A direct comparison between alternative methodologies was conducted using a Wilcoxon Rank Sum Test.

Linear morphological features (mass and SVL) were used to calculate an estimate of individual body condition as suggested by Peig and Green (2009). We first used the *smatr* R-package (Warton *et al.*, 2012) to compute a scaling exponent (derived from a standardized major axis regression). This exponent was then used to calculate the Scaled Mass Index (SMI) for each individual using the following expression:

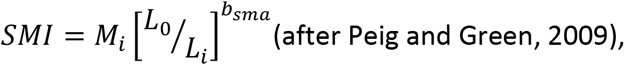

where Mi and Li are the body mass and the linear body measurement of individual *I* respectively; b_sma_ is the scaling exponent estimated by the SMA regression of M on L; L_0_ is an arbitrary value of L (in this case we used the arithmetic mean value of SVL for the study population). We accounted for the significant difference between sexes by computing a sex-specific separate scaling exponent prior to computing the SMI for each individual. This value was used in a regression analysis using hematological and biochemical values as response variables against SMI. We looked for significant correlations using Spearman’s Rank Correlation index, which is more robust when data depart from normal distribution.

### Inter-specific

We compiled morphological and physiological data on Galápagos iguanas from Lewbart and colleagues (2015, 2019). When possible, we used this information to compare estimated values for other species with those recorded for pink iguanas.

## Results

### Intra-specific

We used samples collected in 2019 to assess if different types of i-Stat cartridges (CG8 and Chem8) would produce significantly different results when measuring blood parameters. We compared results for glucose, calcium, sodium, potassium and hematocrit and found no significant differences. The total number of individuals collected between 2019 and 2021 was 42. Of these, one sample escaped while handling it and, although a blood sample had been already collected, we were not able to complete the screening procedure. For this reason, we eliminated it from the analyzed data set. Table 1 shows sampling details of analyzed pink iguana individuals divided by sex and year. As every animal is uniquely tagged, we identified two individuals that were captured in both sampling seasons.

**Table 1.** Number of individual pink iguanas sampled on WV by sex and year (Tot. – total).

|  | **Females** | **Males** | **Tot.** |
| --- | --- | --- | --- |
| *2019* | 7 | 8 | 15 |
| *2021* | 16 | 10 | 26 |
| *Tot.* | 23 | 18 | 41 |

Summary statistic results by sex within pink iguanas are presented in Table 2 and Figures 1, 2 and 3. Morphological characteristics of adult pink iguanas analyzed here are in line with results from other studies (see for example Gentile *et al.*, 2009), with males significantly larger and heavier than females (Wilcoxon Rank Sum Test with continuity correction p-value ≪ 0.01 for SVL and for mass ≪ 0.01). No significant sex effect was found on other variables such as heart rate, respiratory rate or temperature.

**Table 2.** Mean, Standard Deviation (SD), minimum value (Min), and maximum value (Max) of pink iguana morphological features and health parameters by sex (M = male, F = female).

|  | Mean |  | SD |  | Min |  | Max |  |
| --- | --- | --- | --- | --- | --- | --- | --- | --- |
|  | M | F | M | F | M | F | M | F |
| **SVL (cm)** | 47.10 | 41.60 | 4.82 | 4.53 | 37.40 | 33.60 | 60.20 | 51.80 |
| **Mass (Kg)** | 4.99 | 3.76 | 1.25 | 0.82 | 1.50 | 2.20 | 6.50 | 4.87 |
| **HR (bpm)** | 69.83 | 77.82 | 18.33 | 14.80 | 32.00 | 60.00 | 92.00 | 108.00 |
| **RR (brpm)** | 23.05 | 24.52 | 7.88 | 12.55 | 12.00 | 10.00 | 36.00 | 60.00 |
| **T °C** | 31.42 | 31.76 | 5.67 | 5.35 | 21.30 | 22.40 | 39.20 | 40.10 |

**Figure 1.**
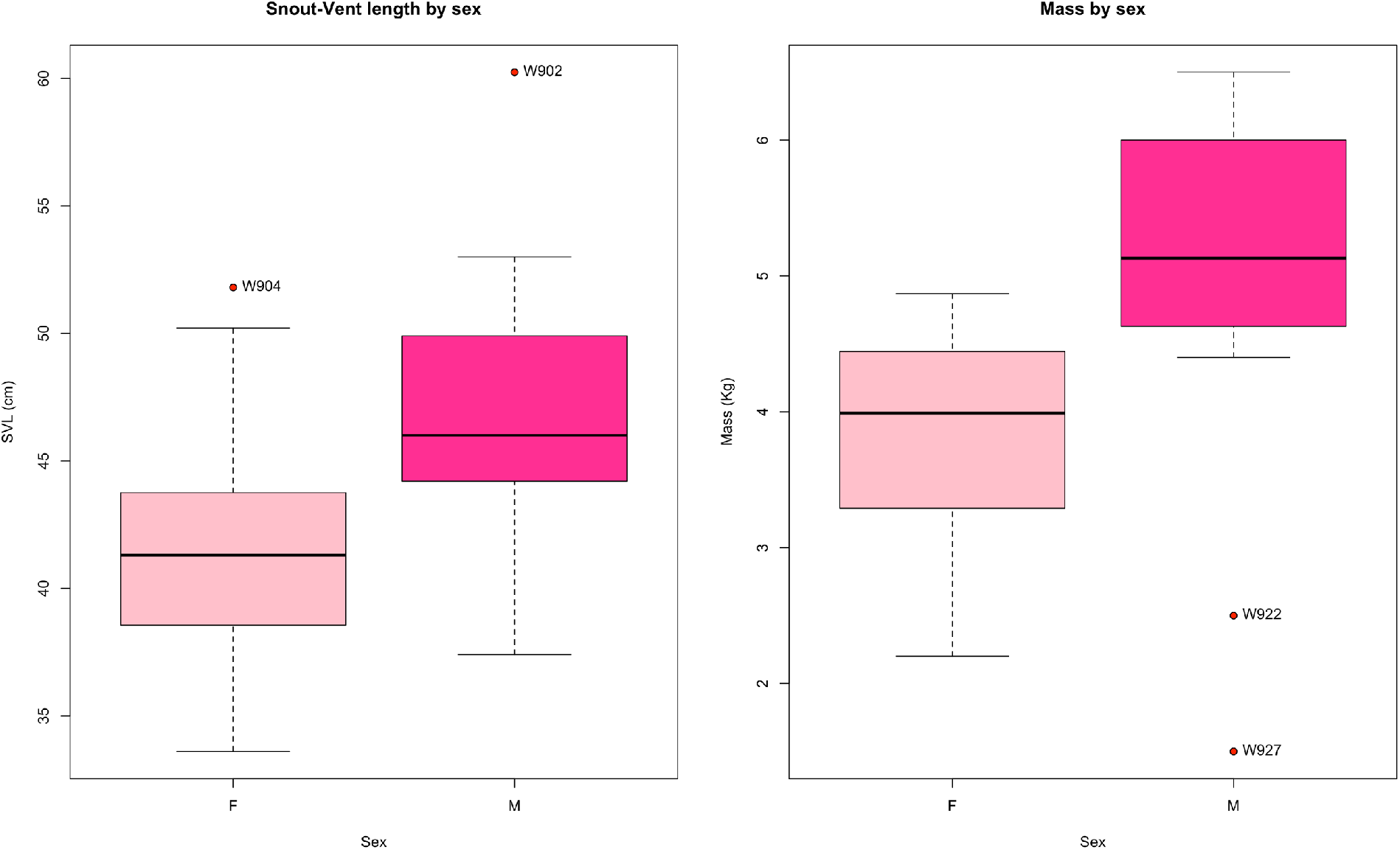
Boxplot representing SVL and body mass data for female (F) and male (M) pink iguanas sampled. These and following boxplots should be interpreted as follows: lower whisker edge represents minimum value; lower box edge represents 25th percentile (Q1); black thick line represents median value; upper box edge represents 75th percentile (Q3); upper whisker edge represents maximum value. Potential outliers are represented by red dots extending beyond the lower and upper whiskers. In all boxplots the text associated with the outliers is the field identifier code of the sampled individuals that are deemed outliers.

**Figure 2.**
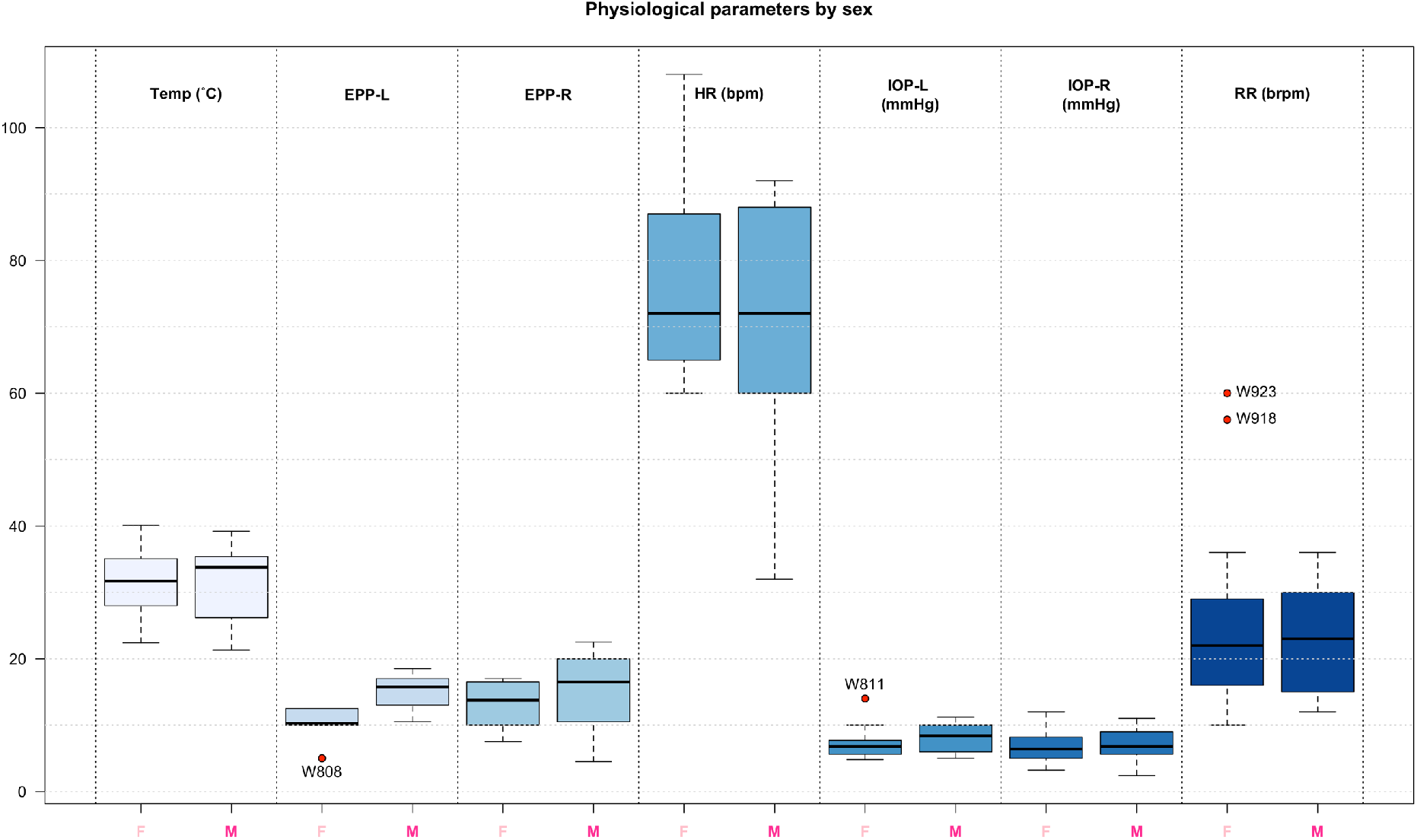
Boxplot comparisons of physiological parameters for female (F) and male (M) pink iguanas.

**Figure 3.**
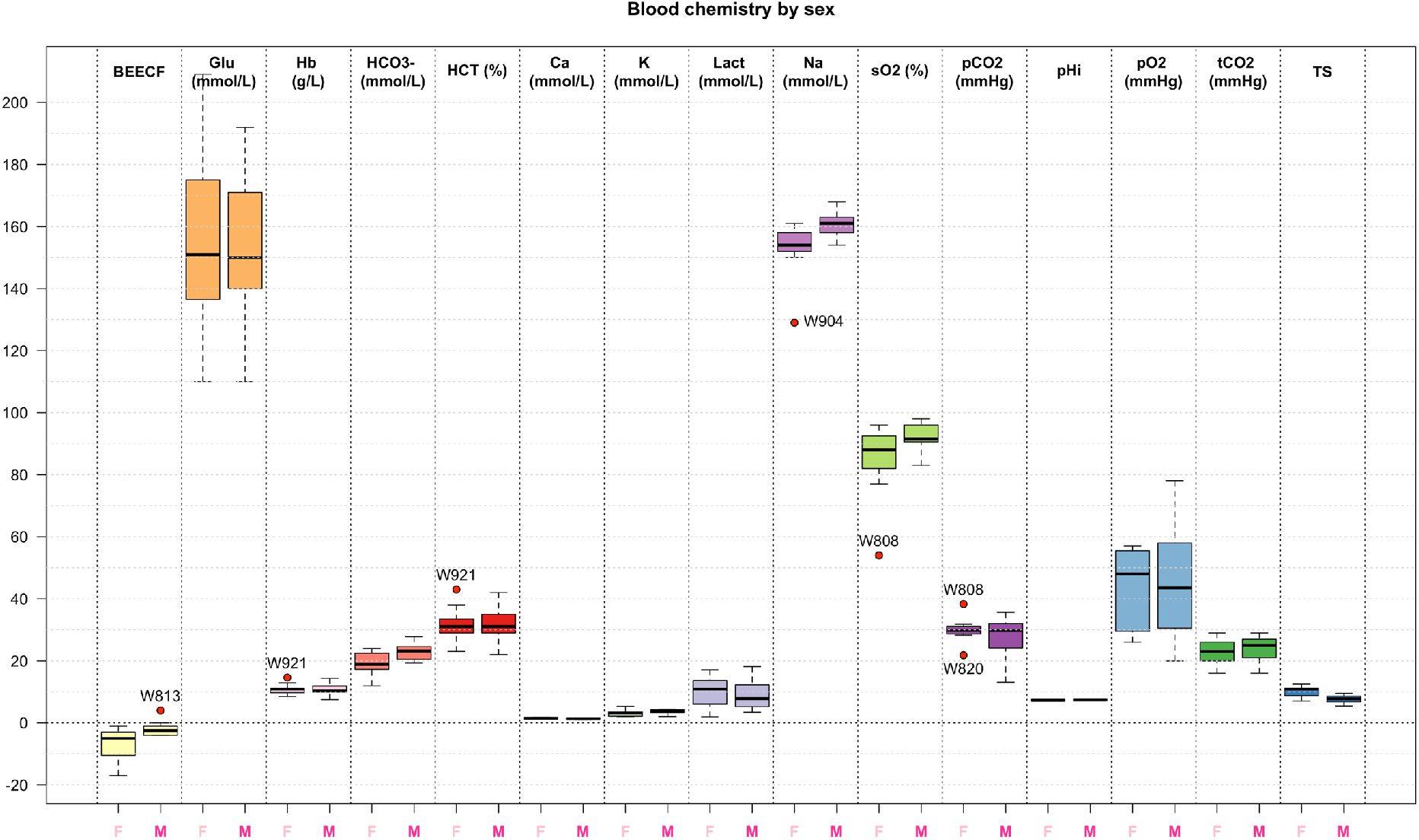
Boxplot comparisons of blood parameters for female (F) and male (M) pink iguanas.

All animals sampled were deemed clinically healthy based upon physical examination with the exception of one 2021 female with a marked dermatitis, likely associated with a heavy mite infestation (Figure 4). Otherwise this animal appeared to be healthy. Two animals were noted to be missing digits or parts of digits, most likely from intraspecific aggression (Figure 5). Almost all iguanas (95%) collected between 2019 and 2021 had ticks (between 9 and 90) on various parts of their body. Ticks were collected and stored in 70% ethanol but remain in Galápagos for identification at a later time. No significant abnormalities were noted on initial examination or on review of the infrared ocular images. A representative image showing the detail captured in the photographs is presented in Figure 6. The intraocular pressure values for right (R) and left (L) eyes were similar and not significantly different after a Wilcoxon rank sum test with continuity correction (mean ± standard deviation IOP_R = 7.04 ± 2.41; mean ± standard deviation IOP_L = 7.50 ± 2.17; p-value = 0.32). Tear production (TP) values for the right eye (mean ± standard deviation TP_R = 14.2 ± 5.22) were not significantly different (p-value = 0.51) for TP values of the left eye (mean ± standard deviation TP_L = 12.9 ± 3.66). No effect of sex was found on any of the general physiological parameter measured and reported (Table 2 and Figures 1 and 2). Table 3 and Figure 3 present summary statistics on all biochemical and blood parameters measured for pink iguanas. We found a significant difference between males and females only in ionized calcium values (Wilcoxon Rank Sum Test with continuity correction p-value = 0.02) and total solids (Wilcoxon Rank Sum Test with continuity correction p-value ≪ 0.01). Figure 7 shows the correlation between different variables for pink iguana individuals.

**Figure 4.**
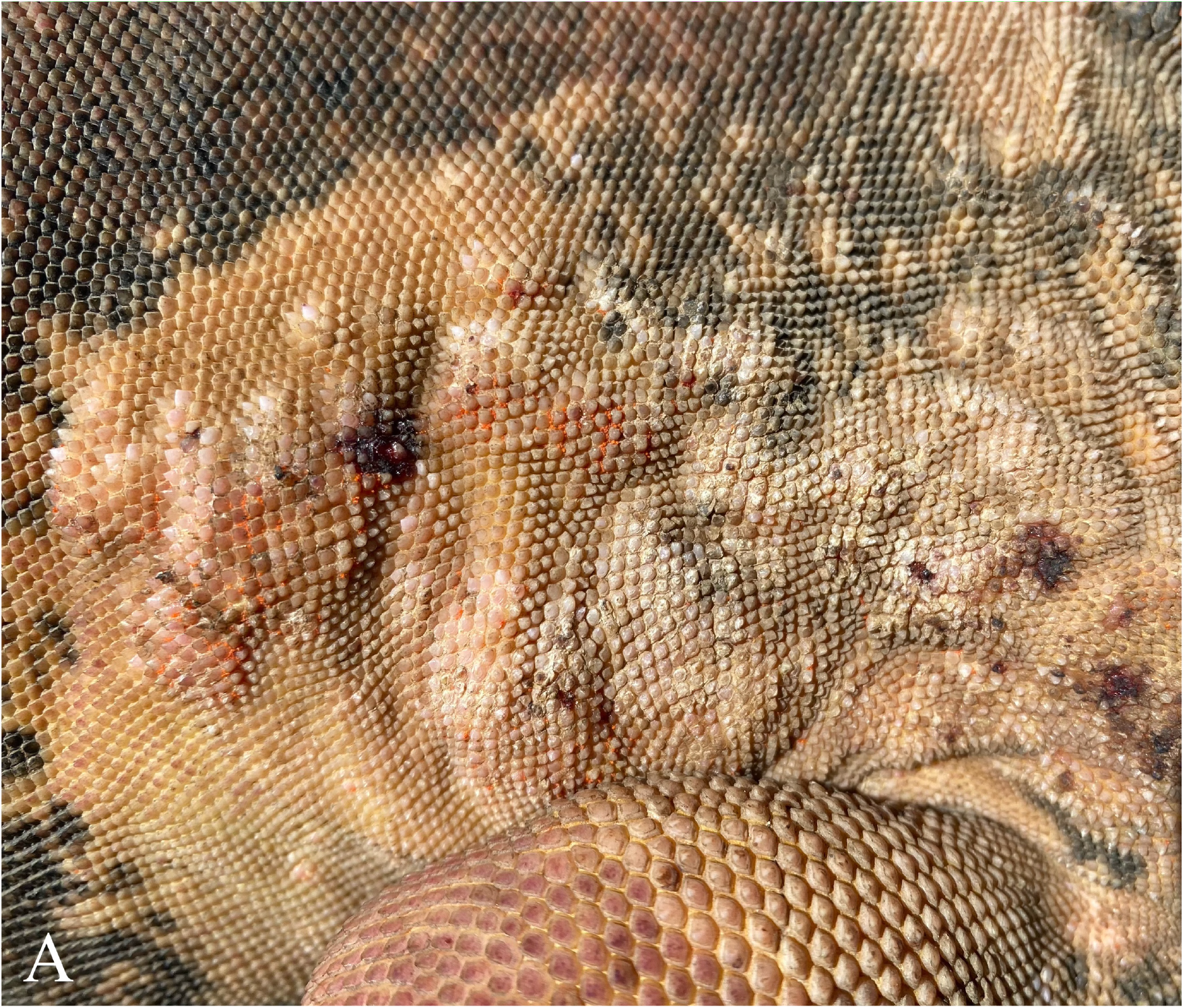
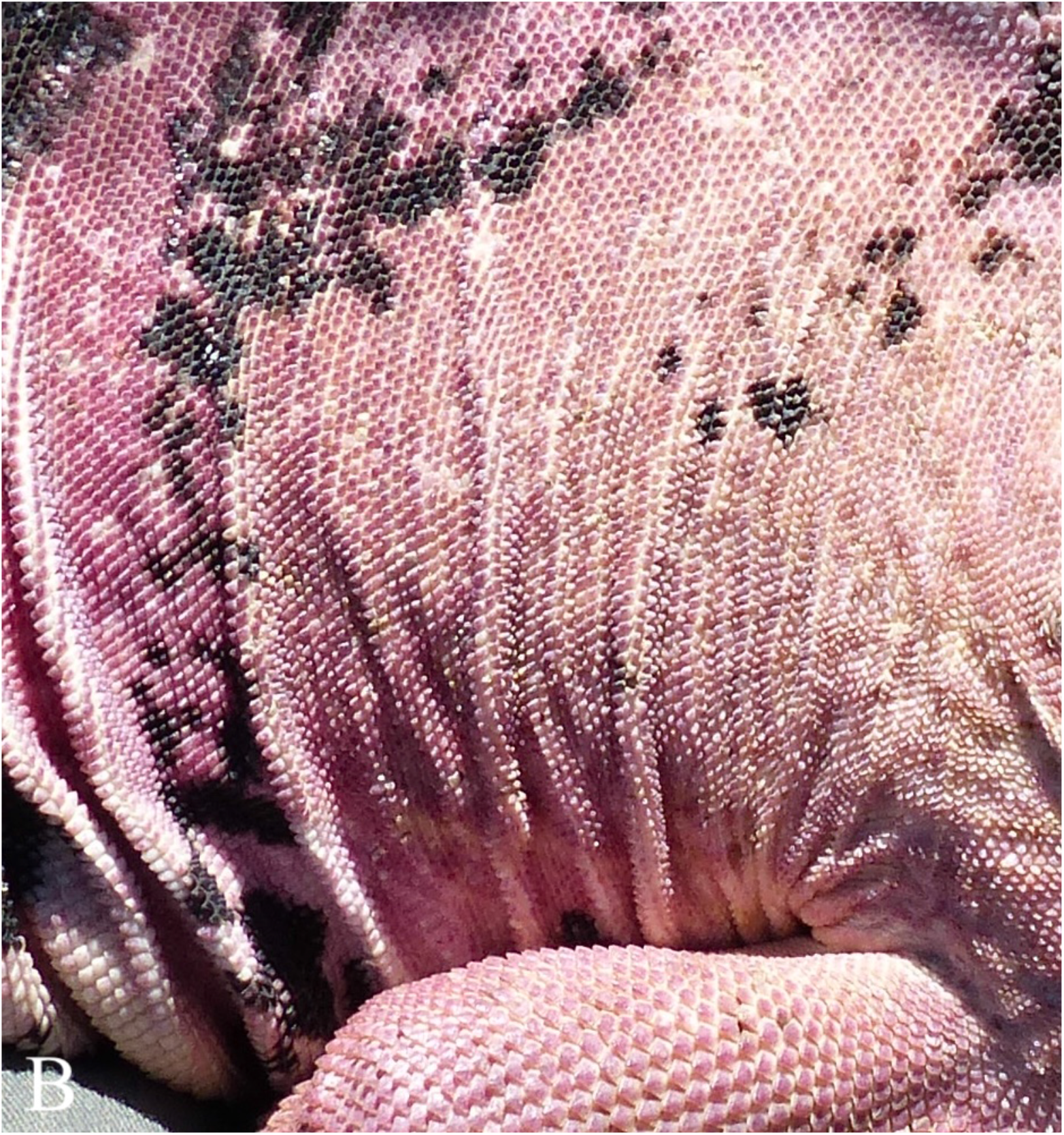
Right side cranial trunk and shoulder of two pink land iguanas (*Conolophus marthae*). A. Adult female with marked raised dermatitis and a heavy mite infestation (clusters of small orange dots). B. Adult female with normal skin and a lack of visible ectoparasites.

**Figure 5.**
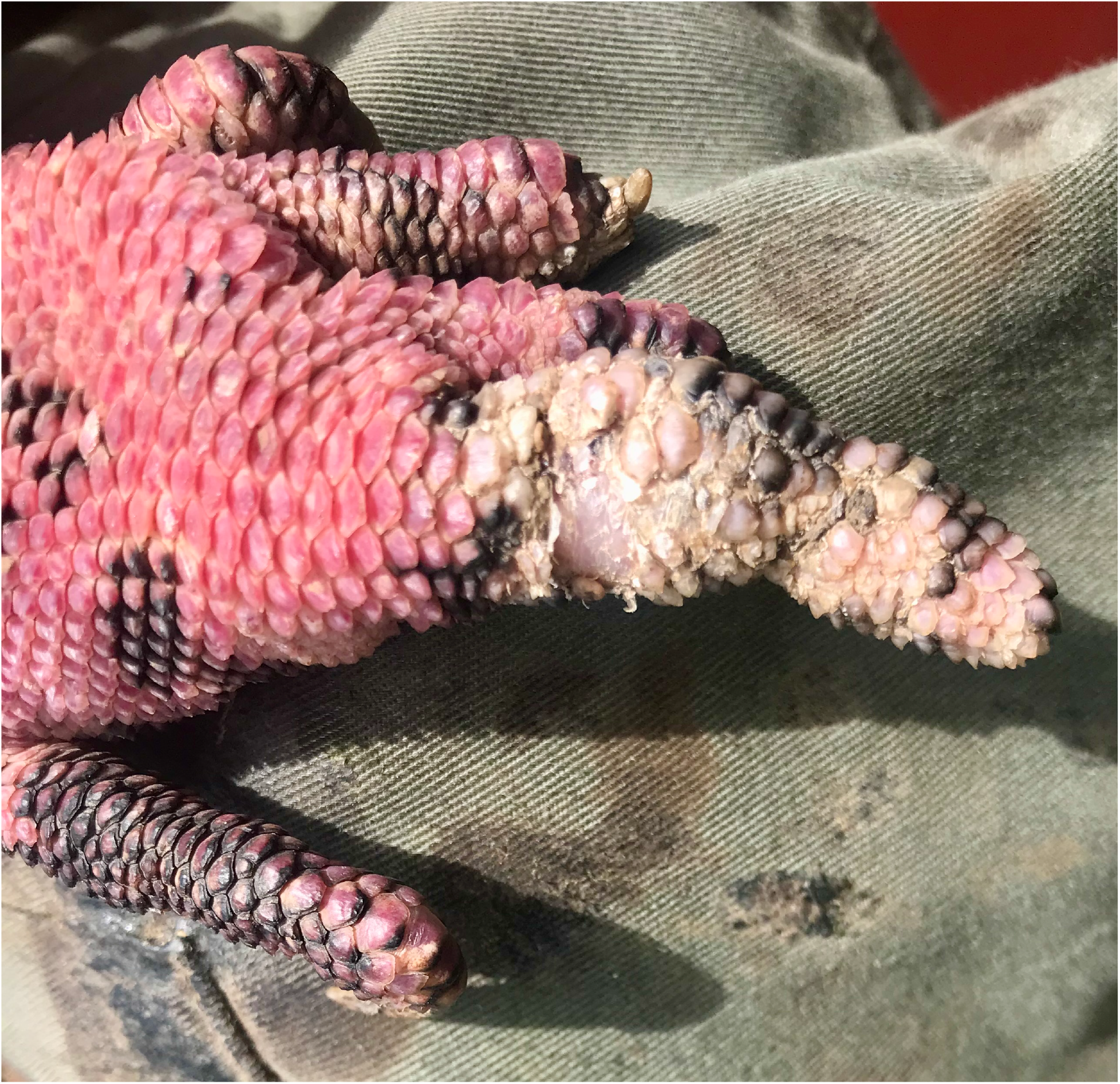
Hind foot of a pink land iguana (*Conolophus marthae*) exhibiting damaged but healed fourth digit.

**Figure 6.**
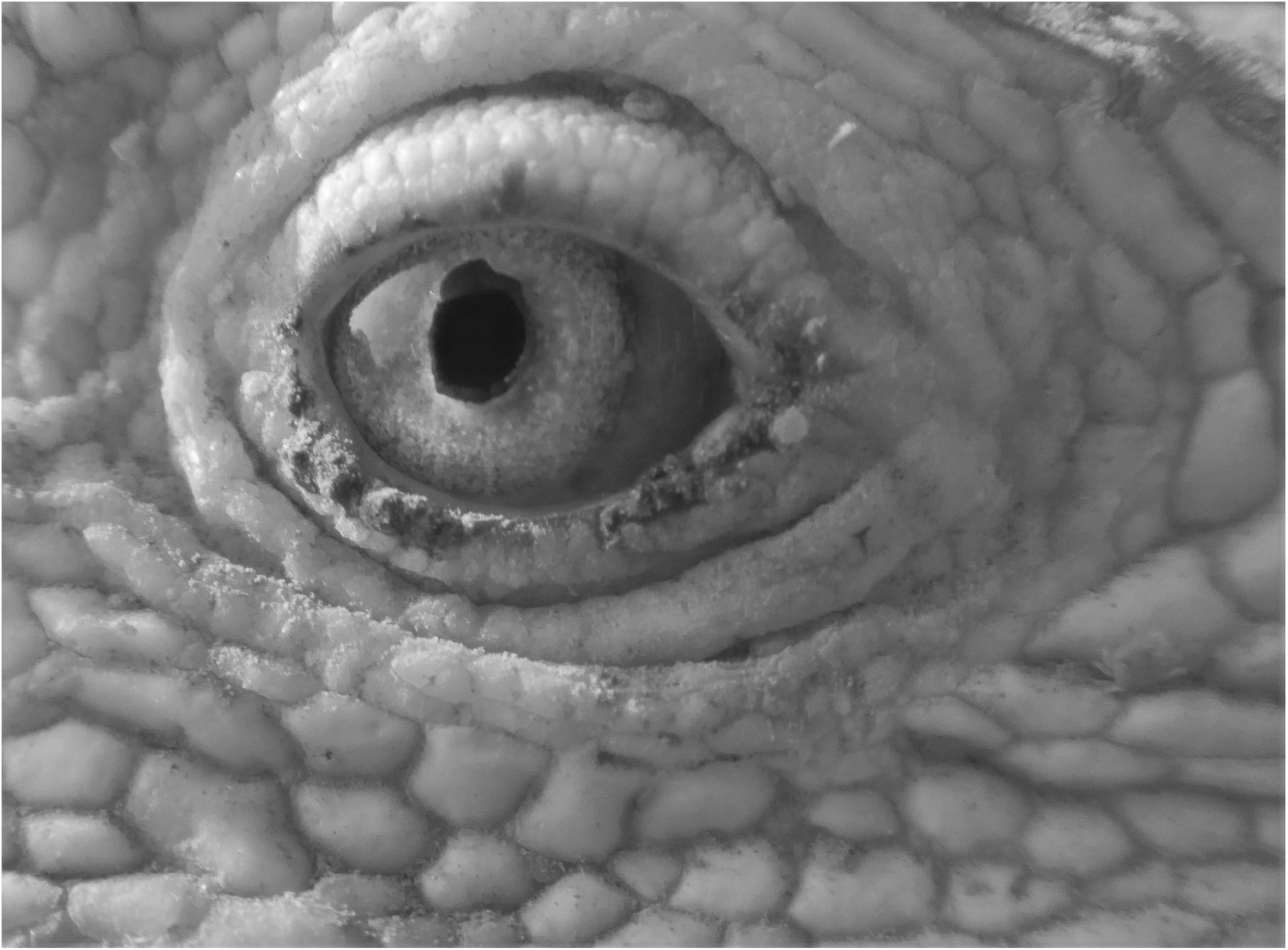
Example of infrared image used to analyze ocular health in pink iguana individuals.

**Figure 7.**
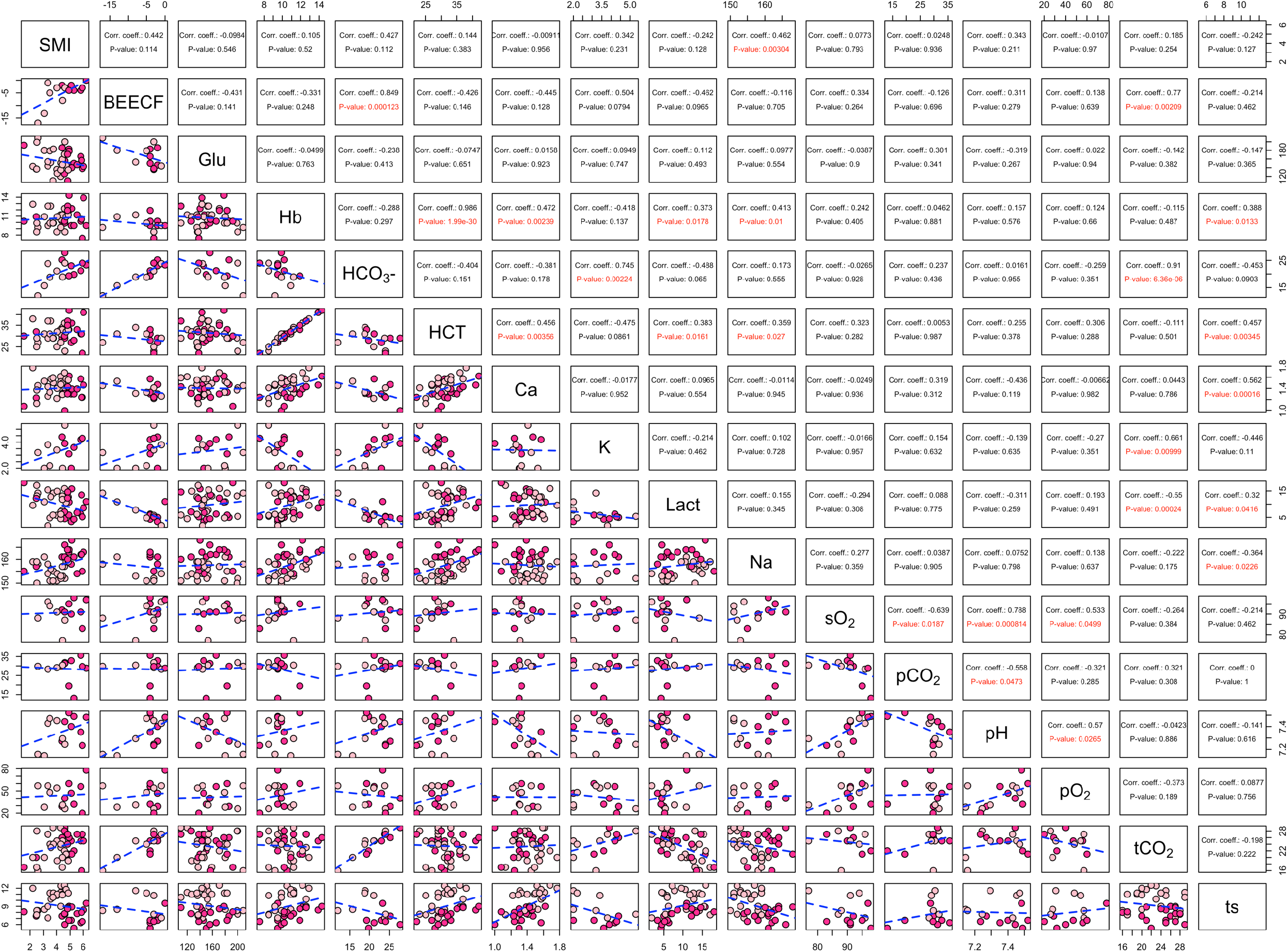
Spearman’s Rank Correlation values estimated on linear regression lines between variables listed across the diagonal. Significant correlations are highlighted in red.

**Table 3.** Mean and Standard Deviation (SD) of blood biochemical parameters measured in *C. marthae* samples and separated according to sex (M = male, F = female). Base excess in extracellular fluid compartment (BEECF); Glucose (Glu); Hemoglobin (Hb); Bicarbonate (HCO_3_^−^); Hematocrit (HCT); ionized calcium (Ca); Potassium (K); Lactate; Sodium (Na); Partial pressure of carbon dioxide (pCO_2_); pH measured at standardized temperature (pH_[37°C]_); Partial pressure of oxygen (pO_2_); Total carbon dioxide (tCO_2_); Total protein solid (TS). Variables highlighted in bold indicate there was a significant difference between males and females.

|  | Mean |  | SD |  |
| --- | --- | --- | --- | --- |
|  | M | F | M | F |
| *BEECF* | -1.87 | -7.14 | 2.74 | 5.89 |
| *Glu (mmol/L)* | 154.47 | 155.21 | 22.41 | 26.19 |
| *Hb (g/L)* | 10.89 | 10.69 | 1.67 | 1.56 |
| *HCO<sub>3</sub><sup>-</sup> (mmol/L)* | 22.95 | 19.18 | 2.83 | 4.32 |
| *HCT (%)* | 32.11 | 31.47 | 5.01 | 4.72 |
| ***Ca (mmol/L)*** | **1.33** | **1.45** | **0.15** | **0.15** |
| *K (mmol/L)* | 3.64 | 3.12 | 0.85 | 1.20 |
| *Lactate (mmol/L)* | 9.18 | 10.25 | 4.45 | 4.35 |
| ***Na (mmol/L)*** | **160.82** | **153.95** | **3.94** | **6.33** |
| *pCO<sub>2</sub> (mmHg)* | 27.57 | 29.92 | 7.47 | 4.88 |
| *pH<sub>[37 °C]</sub>* | 7.39 | 7.31 | 0.10 | 0.13 |
| *pO<sub>2</sub> (mmHg)* | 45.25 | 43.00 | 18.94 | 14.11 |
| *tCO<sub>2</sub> (mmHg)* | 23.88 | 22.91 | 3.78 | 3.98 |
| ***TS (g/L)*** | **7.62** | **10.22** | **1.15** | **1.65** |

### Inter-specific

On average, body temperature and potassium levels did not differ significantly between the four species. For all other comparisons we did find some statistically significant differences (Figures 10 and 11). The respiratory rate of *A. cristatus* turned out to be lower than all species of land iguanas, with that of male *A. cristatus* significantly lower after Bonferroni correction (Figure 11).

**Figure 8.**
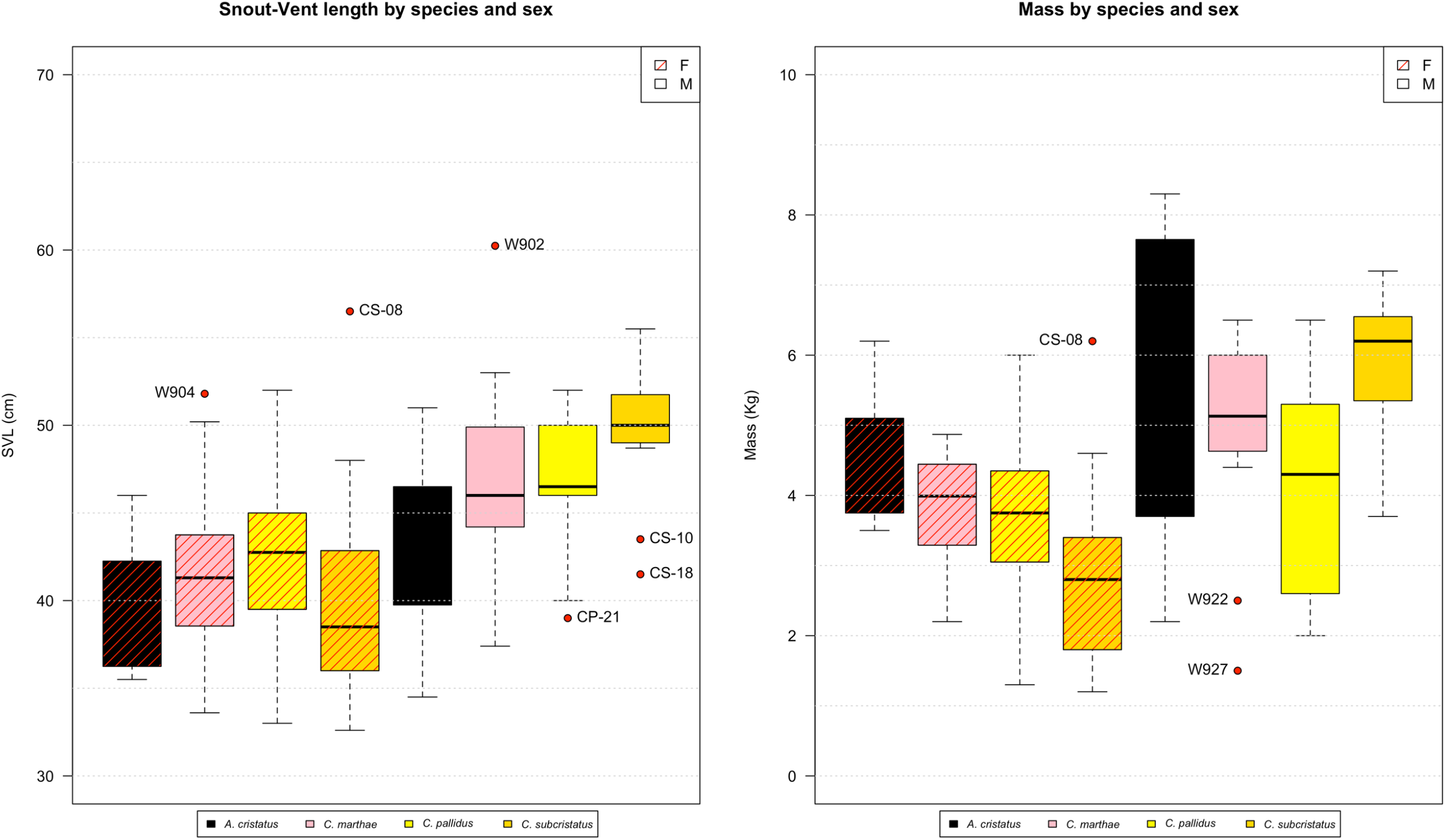
Boxplot comparisons of snout-vent length (SVL) and mass for sampled female (F) and male (M) iguanas by species. Box plot parameters as in Figure 1.

**Figure 9.**
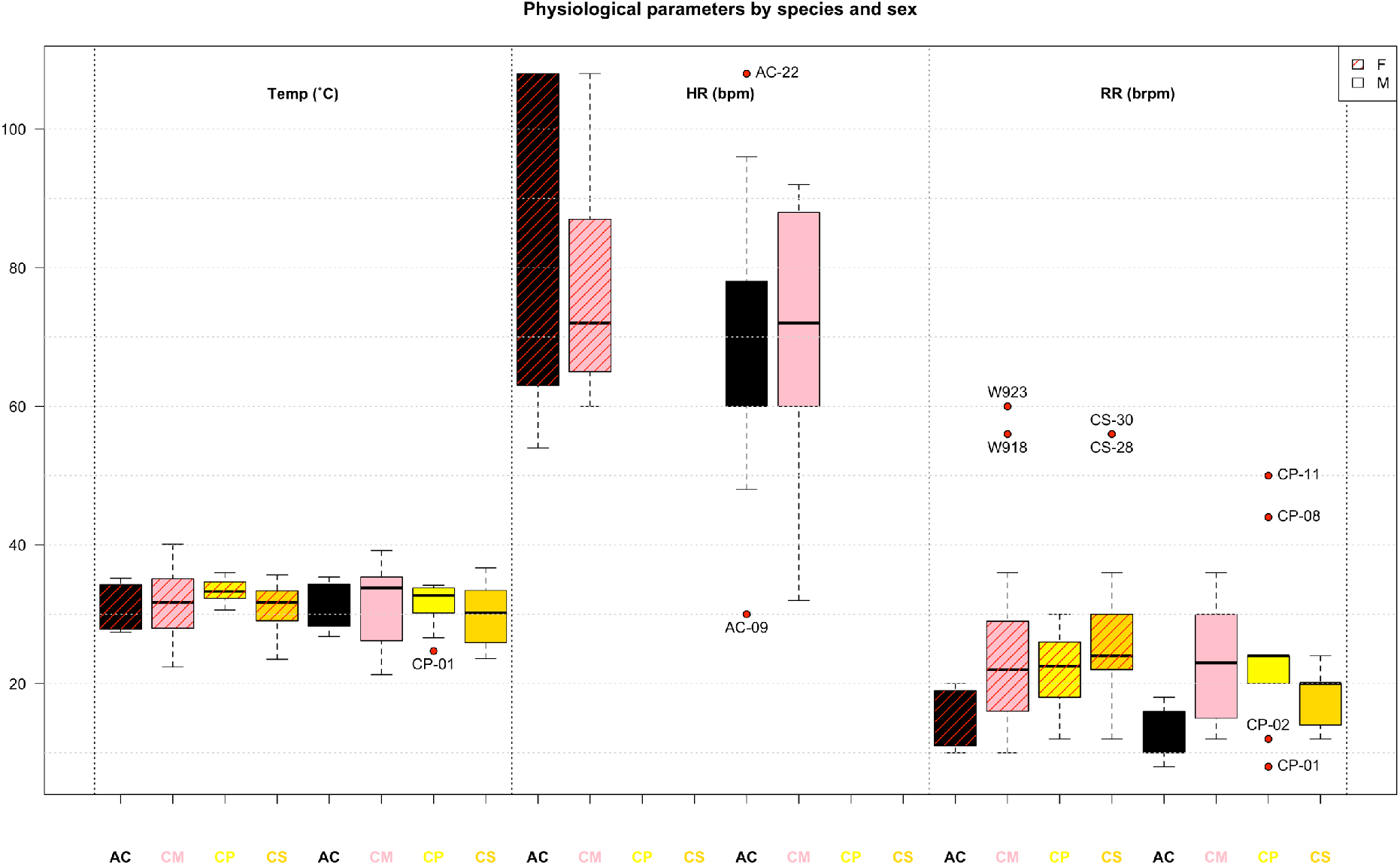
Boxplot comparisons of physiological parameters for sampled female (F) and male (M) iguanas by species. Box plot parameters as in Figure 1.

**Figure 10.**
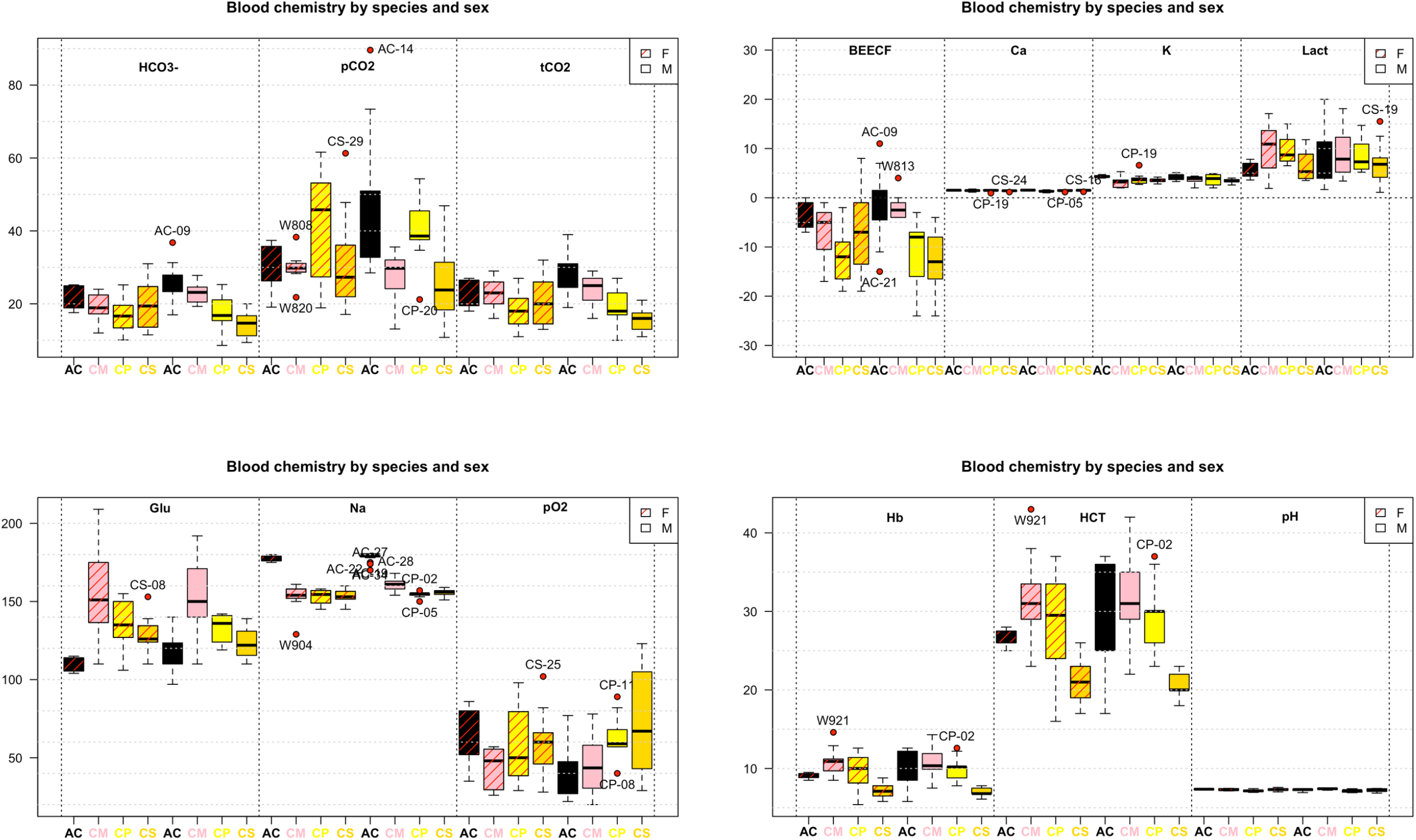
Boxplot comparisons of blood parameters between male and females of different iguana species.

**Figure 11.**
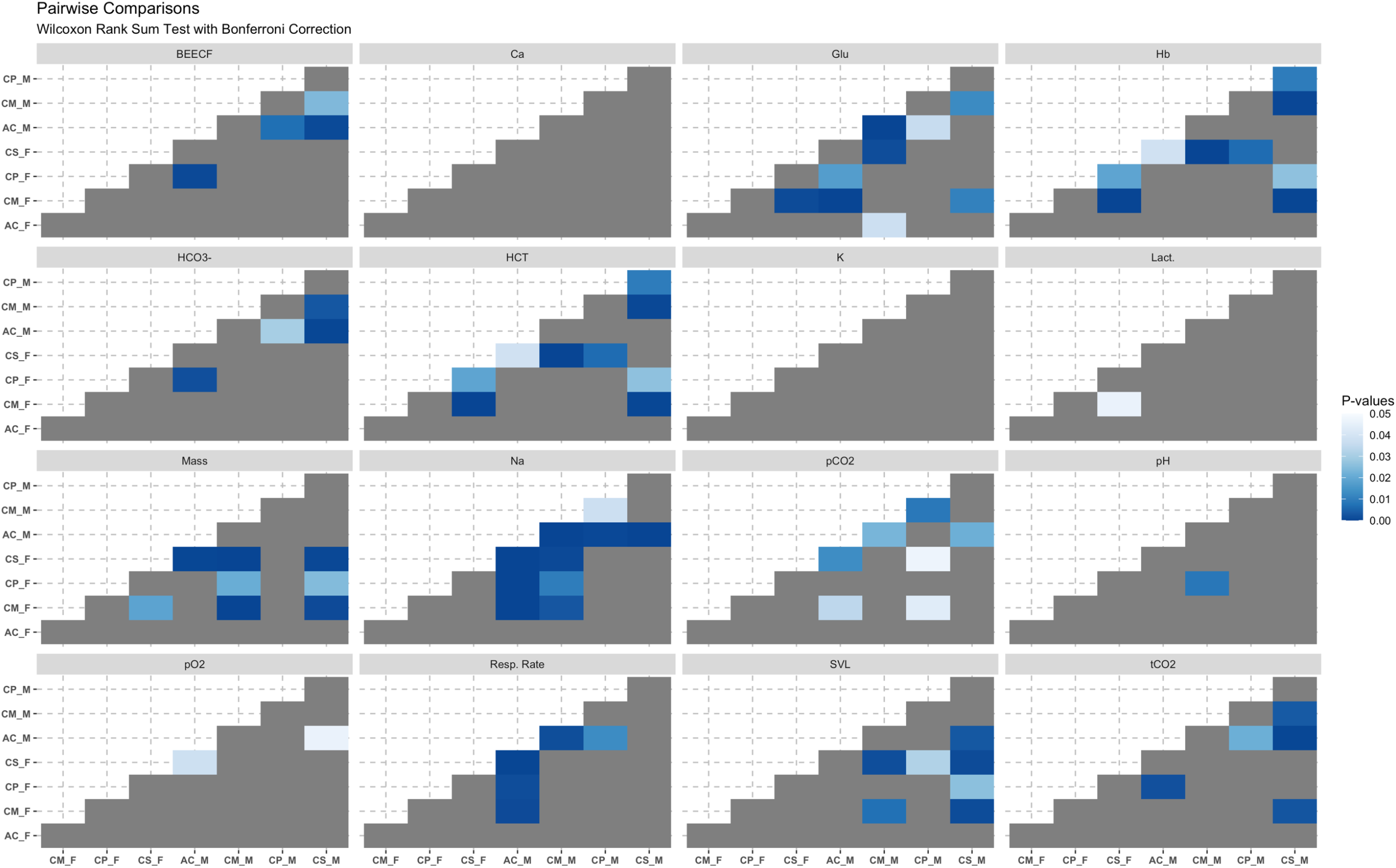
For each estimated parameter we looked for significant differences between species by sex. Each heatmap above shows when values differed significantly between sexes or species. P-values were adjusted for multiple comparisons using Bonferroni correction method. Darker shades of blue indicate p-values closer to 0.

## Discussion

The work of field biologists and veterinarians benefits from having species-specific baseline physiological values from healthy animals for parameters that can be quickly, accurately, and inexpensively measured with commercial equipment. In ectothermic reptiles, such data are especially important, owing to diverse environmental conditions and habitats. Our study provides the first blood gas, biochemistry, and hematology measures in Galápagos pink land iguanas and performs a preliminary comparison of these data with those published for other species of Galápagos iguanas. These results provide a useful baseline for veterinarians and other researchers.

Point-of-care analyzers like the i-STAT may require caution in interpretation, as published studies have found that some blood gas and hematocrit values are not always accurate or reliable with certain non-mammalian species. In rainbow trout (*Onchyrhynchus mykiss*) results varied with temperature and only pH was a uniformly reliable value (Harter *et al.*, 2014). A study in sandbar sharks (*Carcharhinus plumbeus*) determined the i-STAT is not reliable for accurately measuring blood gases (Harter et al., 2015b). The i-STAT did not produce valid sO_2_ or hemoglobin values in the bar-headed goose (*Anser indicus*; Harter *et al.*, 2015). In our case, the blood gas values and pH were fairly consistent between the four species, except for the much higher sodium levels in marine iguanas (Lewbart et al., 2015). This is most likely a result of the dietary and habitat difference between the terrestrial and marine iguanas. Marine iguanas also had higher potassium levels than all three land iguanas. The calcium values were comparable between all four species.

Most of the blood values we recorded for pink land iguanas were similar to those reported in other iguanids (Alberts *et al.*, 1998; Dallwig *et al.*, 2011; Divers *et al.*, 1996; Harr *et al.*, 2001; Klaphake *et al.*, 2017; Lewbart *et al.*, 2019; Maria *et al.*, 2007). For example, the average PCV in green iguanas, *Iguana iguana*, (Harr *et al.*, 2001) was 36.7%, slightly higher than the average found in pink iguanas (31.75%). This value, indeed, is more similar to values reported for the basilisk lizard (*Basiliscus plumifrons*, 31.4%), and much closer to that of *C. subcristatus* (Dallwig *et al.*, 2011; Lewbart et al., 2019). Blood sodium levels reported for green iguanas and basilisk lizards are 160 and 153.5 mmol/L, respectively (Harr et al., 2001; Dallwig *et al.*, 2011), whereas the average sodium levels recorded for *C. marthae, C. subcristatus,* and *C. pallidus* are 156.9, 154.0, and 153.7 mmol/L, respectively. A general and easy way to determine health status and stress is blood glucose (Martinez-Jimenez and Hernandez-Divers, 2007). Basilisk lizards (that were held in cloth bags overnight and sampled 12 hours post-capture) had a fairly high level (203 mg/dL; Dallwig *et al.*, 2011) while *C. subcristatus* and *C. pallidus* sampled almost immediately upon capture had much lower levels (126 and 135 mg/dL, respectively; Lewbart et al., 2019). The pink iguana glucose values, based on the i-STAT, average 154.9 mg/dL, similar to that of captive green iguanas (166 mg/dL for males and 180 mg/dL for females, Harr *et al.*, 2001). Wild Allen Cays rock iguanas (*Cyclura cychlura inornata*) had a mean glucose of 189 mg/dL (James et al., 2006).

On the whole, the pink iguanas we examined were alert, robust, and clinically healthy. This status is supported by all indicators used in this work, with the differences between the marine and land species explained as reflecting differences in their adaptation and ecology. A general healthy status was found in *C. marthae*, despite high tick loads and high levels of hemoparasite (hemogregarine) infection exhibited by this species. In general, even though the highly hemoparasitized population of *C. subcristatus* from Wolf volcano did show significant alteration in some measures of immune function, significant correlation between corticosteroid levels (or body condition index) and the number of ticks or parasitemia were not found in *C. marthae* (Onorati et al., 2017). This supports the hypothesis that, in *C. marthae*, ecto- and hemoparasites can be sufficiently tolerated, without causing disease or inducing clinical signs such as lethargy, open-mouth breathing, weight loss, and dehydration that may be observed in immunocompromised animals (Nardini *et al.*, 2013).

Although *C. marthae* seems to be in good health at present, we recommend that the only existing population be regularly monitored to track possible changes in health status that may be induced by the harsh and changing environment where the species lives. There are currently discussions by a variety of engaged conservation groups about how to best conserve and protect *C. marthae*. Some of these strategies include captive propagation, head-starting of juveniles, and translocation. If any or all of these practices are employed, having baseline data from healthy wild animals will be critically important.

## Acknowledgements

We are indebted to the park rangers of the Galápagos National Park for their invaluable support and friendship. This work is part of a long-term institutional agreement between the University Tor Vergata and the Galápagos National Park Directorate, aimed at the conservation of Galápagos iguanas. G.C. was supported by a Post-Doctoral Research Fellowship from the San Diego Zoo Wildlife Alliance funded by a donation from the Kenneth and Anne Griffin Foundation. G.G. was supported by grants from the International Iguana Foundation and from Friends of Galápagos, Switzerland. GAL thanks Diego Páez-Rosas, Juan Pablo Muñoz-Pérez, Carlos Mena, Stephen Walsh, and the Galápagos Science Center for their assistance and support.

